# Stimulus identity rather than emotion drives EEG classification on the FACED dataset

**DOI:** 10.64898/2026.06.12.731889

**Authors:** Moritz Gerster, Elizaveta Sirotina, Anton Orlovskii, Alexandra Hertz, Juliette Champaud, Domenico Guarino, Silvia Tulli

**Affiliations:** Human Augmented Brain System (HABS), Paris, 75009, France; Institute of Intelligent Systems and Robotics (ISIR) - CNRS - Sorbonne University

## Abstract

Reliable benchmark datasets are critical for advancing EEG-based emotion recognition. The Finer-grained Affective Computing EEG Dataset (FACED) is the largest publicly available EEG emotion dataset (123 subjects, nine emotion categories) and a widely adopted benchmark. We demonstrate that both intra-subject and cross-subject classification on FACED primarily reflects stimulus identity rather than emotion. Using a linear classifier (LinearSVC) and a deep learning model (CLISA), we show that (1) classification performance is comparable for trials where subjects reported feeling versus not feeling the assigned emotion; (2) accuracy drops when stimulus-assigned labels are replaced with individual self-reports; and (3) accuracy increases when reducing to one video per emotion despite discarding two-thirds of the data. These results reflect three design choices in FACED: few stimuli per category, stimulus-assigned labels, and within-video temporal splits for cross-validation. Together, these make the dataset susceptible to temporal autocorrelation and stimulus-identity confounds. To guide future work, we propose five recommendations — spanning stimulus diversity, temporal independence, and label validation — for emotion-decoding study designs that mitigate these confounds.

## Introduction

Electroencephalography (EEG)-based emotion recognition has attracted growing interest for its potential to objectively measure affective states^1^. Progress in the field depends on benchmark datasets, yet the most established ones — SEED (15 subjects, 3 categories)^2^ and DEAP (32 subjects, 2 dimensions)^3^ — are limited in sample size and emotional granularity. FACED^4^ addresses this gap with a substantial increase in sample size and nine discrete emotion categories, making it, to our knowledge, the largest publicly available EEG emotion dataset to date. Since its release, FACED has been adopted as a standard benchmark: it is included in TorchEEG^5^ and Meta’s NeuralBench-EEG v1.0^6^, and multiple studies have used it to evaluate novel deep learning architectures for cross-subject emotion recognition.

However, classification of neurophysiological signals is vulnerable to confounds that inflate accuracy without reflecting the cognitive or affective state of interest. Brouwer et al.^7^ formulated six recommendations to avoid common pitfalls when using neurophysiological signals for mental state estimation, against which Alarcão & Fonseca^8^ evaluated existing affective computing studies. More recently, Kilgallen et al.^9^ formalized the repeated-stimulus confound in EEG decoding, showing that when a classifier is trained and evaluated on responses to the same stimuli, stimulus identity becomes a confounder for accuracy. For instance, a model may achieve high cross-subject accuracy simply by recognizing the shared neural or artifactual response to a particular clip, rather than learning a generalizable emotion representation.

We show that FACED is vulnerable to these confounds: design choices that conflate stimulus identity with the target emotion label allow classifiers to exploit confounds rather than learn emotion representations. Using both a classical and a deep learning pipeline, we demonstrate that classification on FACED primarily reflects stimulus identity, and propose recommendations for future study designs.

## Methods

### Ethics statement

This study is a secondary analysis of publicly available de-identified data. The original study was approved by the Ethics Committee of Tsinghua University (THU201906) and informed consent was obtained from all subjects^4^.

### Dataset

The FACED dataset^4^ contains 32-channel EEG from 123 subjects who each watched 28 video clips covering nine emotion categories (anger, disgust, fear, sadness, neutral, amusement, inspiration, joy, tenderness). Each non-neutral category comprised three clips; neutral comprised four. The last 30 seconds of each clip served as the analysis epoch. Two mastoid channels were excluded, leaving 30 channels. After each clip, subjects rated their experience on the eight non-neutral emotion items (0–7 scale); however, the original classification pipeline did not use these self-reports and instead assigned each trial the emotion label of its video category.

### Feature extraction and normalization

We used the official pre-computed differential entropy (DE) features published with the dataset rather than re-deriving them from the raw EEG. These comprise five frequency bands (delta: 1–4 Hz, theta: 4–8 Hz, alpha: 8–14 Hz, beta: 14–30 Hz, gamma:30–47 Hz) in non-overlapping 1-second windows, yielding 30 windows per trial and 150 features per window (5 bands× 30 channels). Following Chen et al.^4^, we applied running normalization with exponential decay (rate = 0.990) followed by linear dynamical system (LDS) smoothing per subject.

### Part 1: Intra-subject decoding

#### 1a. Intra-subject baseline

A 10-fold cross-validation split each 30-second video epoch into held-out segments of 3 consecutive seconds (Fig. 1a). All 123 subjects were pooled; the same subject’s data from the same video appeared in both training and test sets, separated only in time. A LinearSVC was trained on the remaining 27 seconds. The hyperparameter *C* was selected from 12 logarithmically spaced values (10^*−*5^ to 10^0.5^) by accuracy on the held-out segment, following the original methodology. This constitutes hyperparameter selection on the test set, which optimistically biases accuracy; we retained it for faithful replication. Accuracy was computed per subject and then averaged.

**Figure 1.**
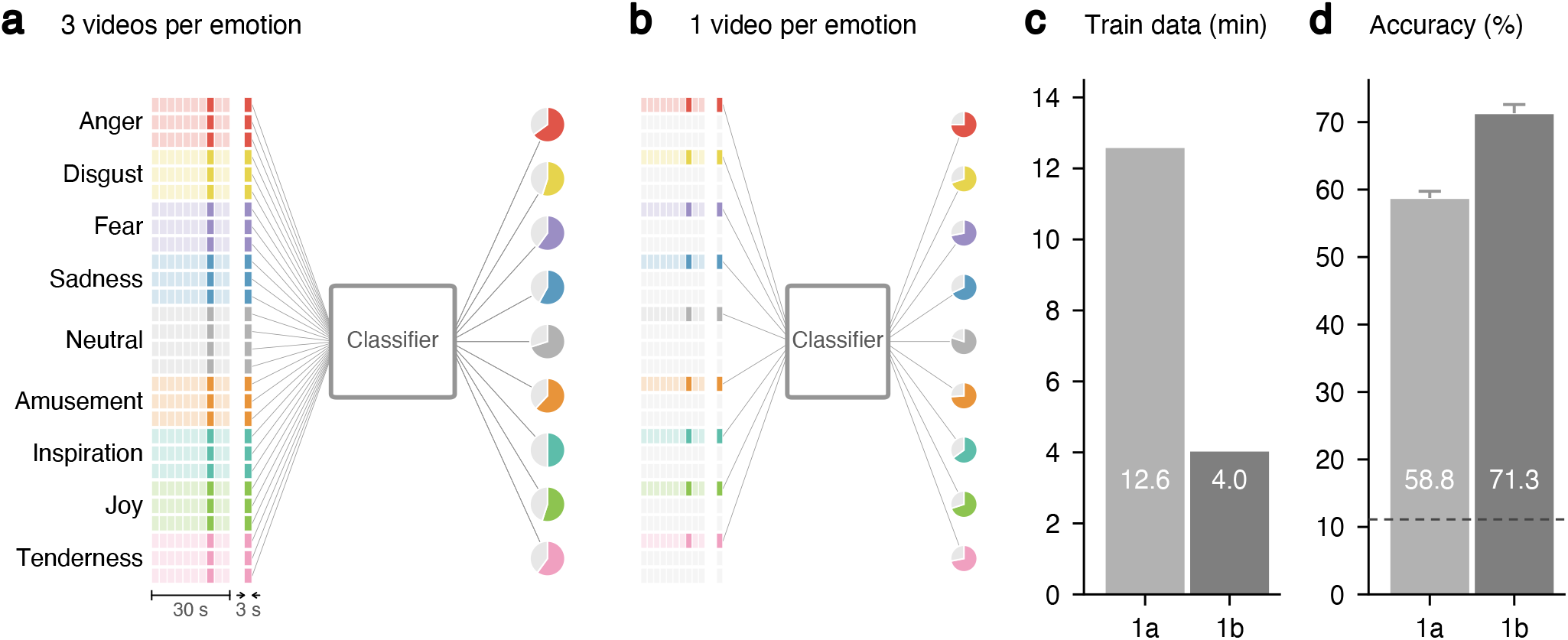
Intra-subject 9-class emotion decoding (Experiments 1a and 1b). **(a)** Baseline pipeline: 28 videos (9 emotions) are each split into 10 folds (27 s train, 3 s test). Test segments are classified by an SVC into 9 categories; pie charts illustrate per-class accuracy (illustrative only). **(b)** Single video per emotion (Experiment 1b): only the first video per emotion is retained (colored); removed videos shown in gray. **(c)** Training data: 12.6 min (1a) vs. 4.0 min (1b). **(d)** Mean accuracy (±SEM). Despite using only one-third of the data, accuracy increases from 58.8% to 71.3%. Dashed line: chance (11.1%).

#### 1b. Single video per emotion

We repeated experiment 1a but restricted the stimulus set to one video per emotion category (9 videos total; Fig. 1b), discarding two-thirds of the data. High accuracy under this design would indicate that the classifier identifies individual video segments via temporal autocorrelation rather than learning shared emotion patterns across multiple stimuli.

### Part 2: Cross-subject decoding

#### 2a. Cross-subject baseline

A LinearSVC was trained and evaluated via 10-fold cross-validation over subjects (nine folds of 12, one of 15 subjects; Fig. 2a). *C* was selected by accuracy on the held-out fold (same optimistic bias as 1a; corrected in 2d). Accuracy was computed per subject, then averaged.

**Figure 2.**
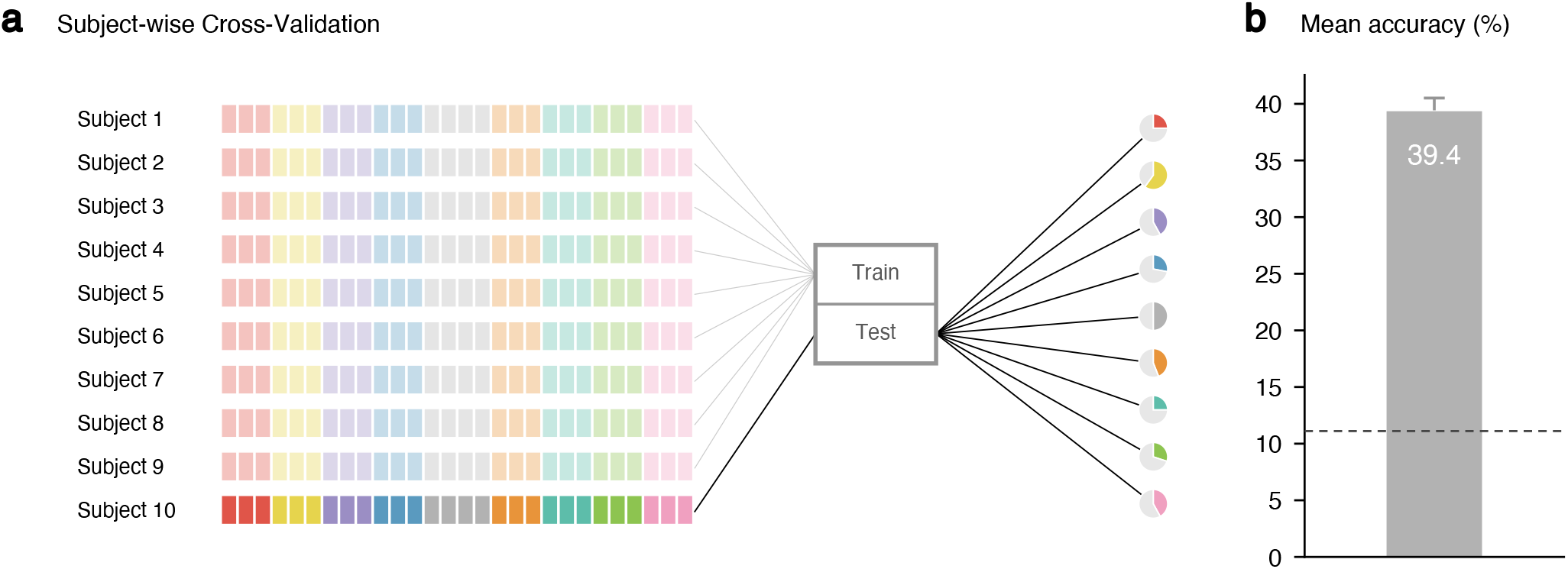
Cross-subject 9-class emotion decoding pipeline (Experiment 2a). **(a)** Subject-wise cross-validation: 123 subjects are split into 10 folds. Nine folds (faded colors) serve as training data and one fold (saturated colors) as test data. Each subject contributes 28 video trials. The classifier is split into Train and Test stages; test-subject predictions are evaluated per emotion category (pie charts are illustrative only). **(b)** Mean accuracy across subjects (*±* SEM). Dashed line indicates chance level (11.1%).

#### 2b. Concordance analysis

After each video, subjects rated the eight non-neutral emotion items on a 0–7 scale. Here, the dominant emotion for a given subject–video pair was defined as the item with the highest rating. To test whether classification performance depends on subjective experience, we partitioned the baseline classifier’s (2a) predictions on non-neutral trials into concordant and discordant subsets (Fig. 3a). A trial was *concordant* if the subject’s dominant self-reported emotion matched the video’s assigned category, and *discordant* otherwise. Neutral videos were excluded because no corresponding self-report item exists. Since the classifier still selects among all nine classes, chance remains at 11.1%.

**Figure 3.**
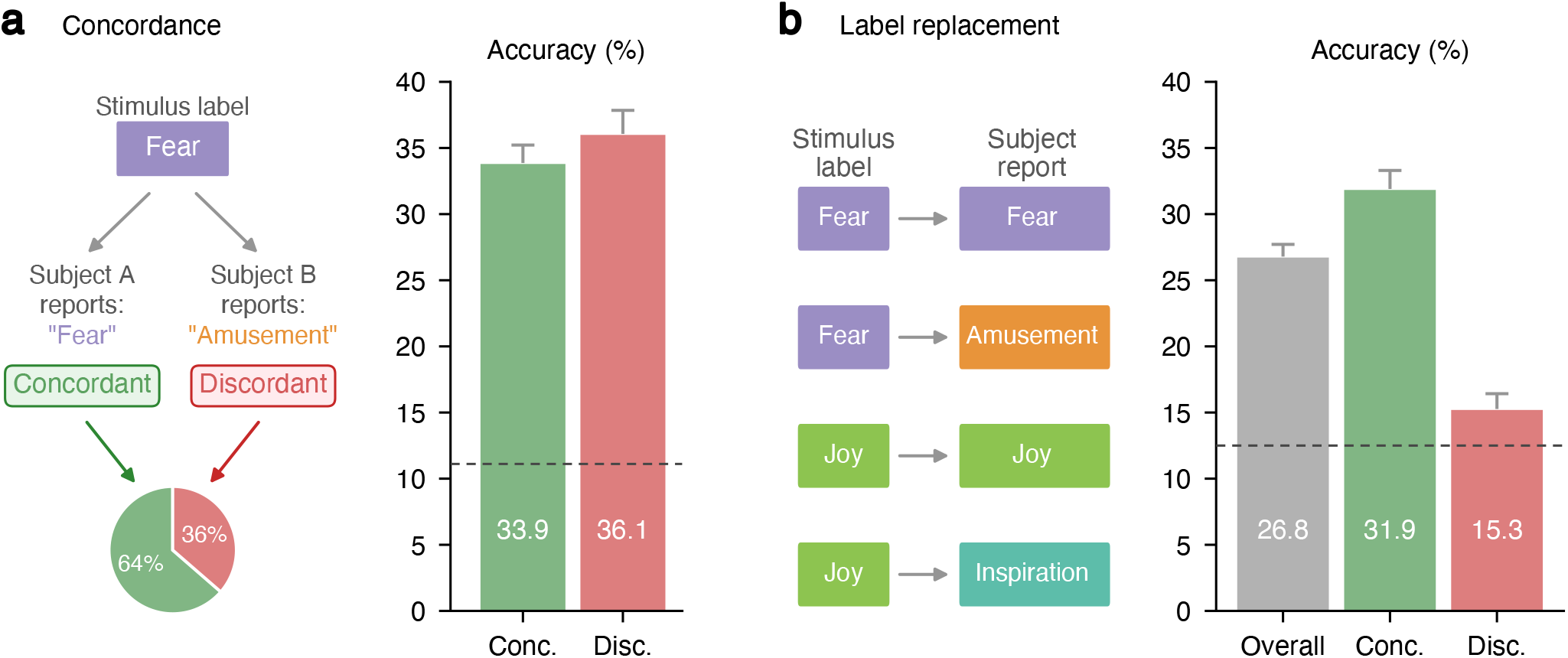
Concordance analysis and subjective labels (Experiments 2b and 2c). **(a)** Concordance definition: a trial is *concordant* if the subject’s dominant self-reported emotion matches the video’s assigned label, and *discordant* otherwise. Pie chart shows the ratio across all non-neutral trials (64% concordant, 36% discordant). Bar plot: mean accuracy (±SEM) of the baseline classifier (2a) split by concordance. Performance is comparable regardless of subjective agreement. **(b)** Label replacement (Experiment 2c): stimulus-assigned labels are replaced with each subject’s self-reported dominant emotion. Bar plot: overall accuracy with subjective labels, and concordance split. On discordant trials, accuracy drops closer to chance (12.5%), indicating the classifier largely fails to predict self-reported emotions that differ from the stimulus label.

#### 2c. Subjective labels

We repeated the cross-subject baseline training but replaced stimulus-assigned labels with subjective labels (Fig. 3b): for each subject and video, the emotion rated highest in that subject’s self-report served as the label. Because no neutral self-report item exists, subjective labels span eight emotions; neutral videos receive the highest-rated non-neutral emotion. The classifier thus has eight output classes (chance: 12.5%). We additionally split the subjective classifier’s predictions into concordant and discordant subsets (as in 2b). Because class distributions within the discordant subset are unbalanced, we report balanced accuracy for the concordance split of 2c; all other conditions use standard accuracy to match the original methodology.

#### 2d. Single video per emotion

To disentangle stimulus identity from emotion representation, we reduced the stimulus set from three videos to one per category (Fig. 4a), discarding two-thirds of the training data but eliminating within-class stimulus variance. We repeated the cross-subject classification from 2a. Hyperparameter *C* was selected via nested 5-fold cross-validation on the training set, eliminating the optimistic bias present in 2a.

**Figure 4.**
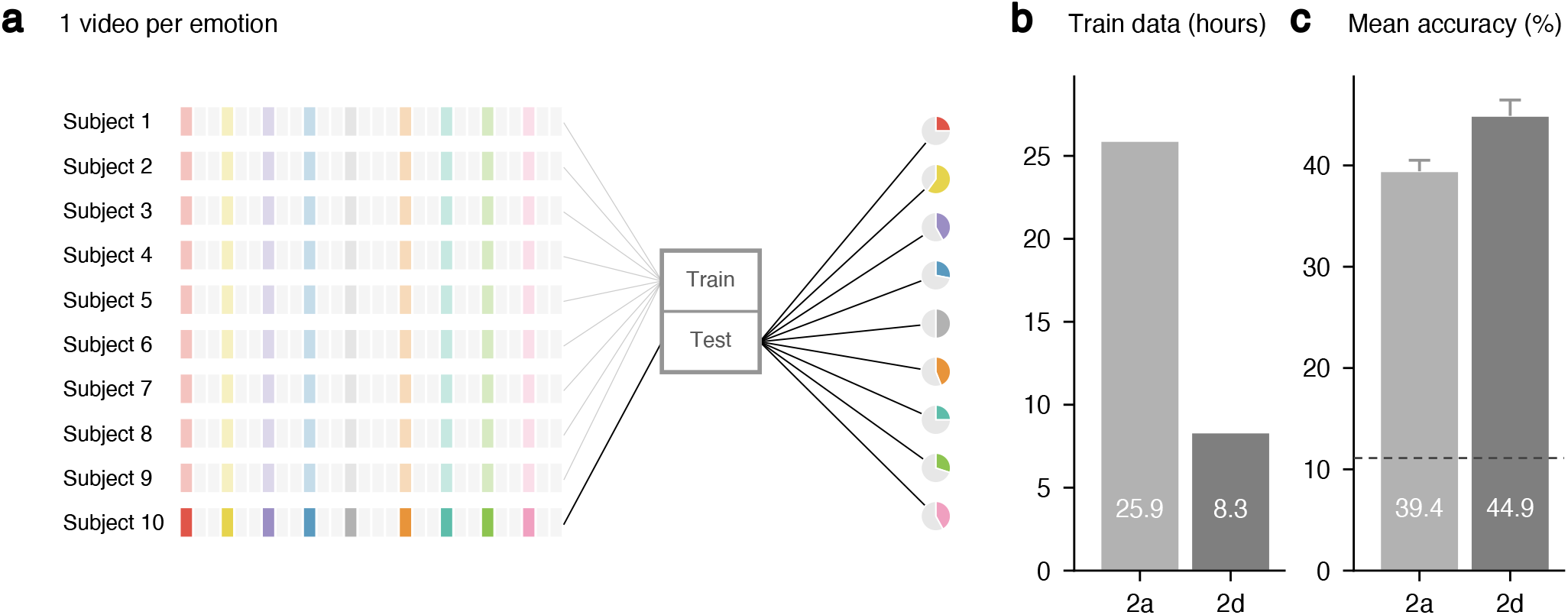
Single video per emotion, cross-subject (Experiment 2d). **(a)** The pipeline from Experiment 2a is repeated but only the first video per emotion is retained (colored bars); removed videos are shown in light gray. Training data is reduced by two-thirds. **(b)** Training data comparison: 25.9 h with all videos (2a) vs. 8.3 h with one video per emotion (2d). **(c)** Mean accuracy (± SEM). Despite the reduced training set, accuracy increases from 39.4% to 44.9%, consistent with the removal of within-class stimulus variance. Dashed line indicates chance level (11.1%).

#### 2e. Deep learning model (CLISA)

To test whether the confound pattern persists beyond linear classifiers, we repeated experiments 2a–2d using CLISA (Contrastive Learning for Inter-Subject Alignment)^10^ (Fig. 5), a deep learning model used by Chen et al. (2023) for dataset validation. CLISA uses contrastive learning to align EEG representations across subjects and was reported to achieve 42.35±1.00% nine-class cross-subject accuracy on the same DE features. We used the published CLISA implementation with default hyperparameters^4^; the held-out fold served for early stopping.

**Figure 5.**
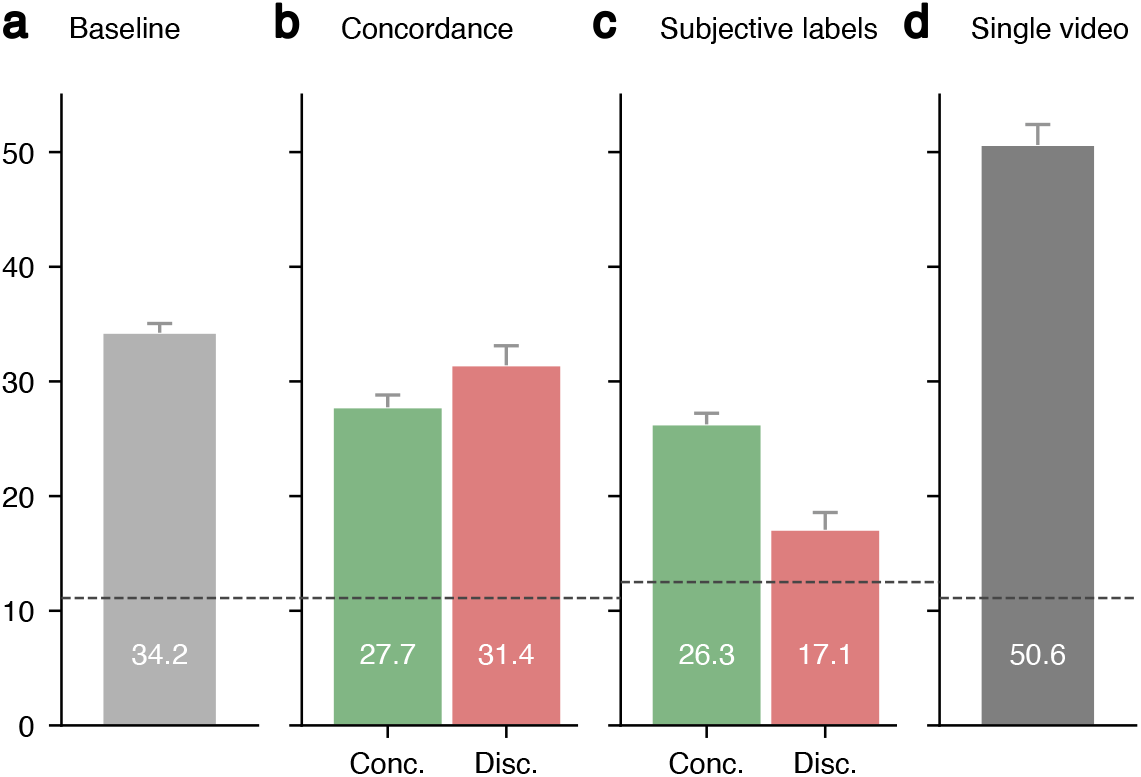
CLISA deep learning results (Experiment 2e). The confound pattern observed with LinearSVC is reproduced with a deep learning model. **(a)** Baseline cross-subject accuracy. **(b)** Concordance split: concordant and discordant trials yield comparable accuracy. **(c)** Subjective labels: accuracy drops on discordant trials (chance: 12.5%). **(d)** Single video per emotion: accuracy increases despite reduced training data. Dashed line indicates chance level (11.1%; 12.5% for subjective labels).

## Results

All accuracies are reported as mean ± standard error of the mean (SEM) across subjects and are summarized in Table 1. Chance level for nine-class classification is 11.1% (12.5% for the eight-class subjective-label analyses).

**Table 1.** Decoding accuracy (mean±SEM across subjects). Chance level = 11.1% (nine classes), except Experiment 2c which uses eight subjective-label classes (chance = 12.5%).

| Exp. | Description | Accuracy (%) | Notes |
| --- | --- | --- | --- |
| *Part 1: Intra-subject decoding* |  |  |  |
| 1a | Baseline (3 videos/emotion) | $58.8 \pm 1.0$ | cf. $51.1 \pm 0.9$ (Chen et al., 2023) |
| 1b | Restricted (1 video/emotion) | $71.3 \pm 1.3$ | ↑ despite $1/3$ data |
| *Part 2: Cross-subject decoding (LinearSVC)* |  |  |  |
| 2a | Baseline (3 videos/emotion) | $39.4 \pm 1.1$ | cf. $35.2 \pm 1.0$ (Chen et al., 2023) |
| 2b | Baseline, concordant trials | $33.9 \pm 1.4$ | No concordance effect |
| | Baseline, discordant trials | $36.0 \pm 1.8$ | |
| 2c | Subjective labels (all trials) | $26.8 \pm 0.9$ | |
| | Subjective labels, concordant | $31.9 \pm 1.4$ | Strong concordance effect |
| | Subjective labels, discordant | $15.3 \pm 1.2$ | |
| 2d | Restricted (1 video/emotion) | $44.9 \pm 1.6$ | ↑ despite $1/3$ data |
| *Part 2: Cross-subject decoding (CLISA)* |  |  |  |
| 2e | Baseline | $34.2 \pm 0.8$ | cf. $39.4 \pm 1.1$ (2a) |
| | Baseline, concordant trials | $27.7 \pm 1.1$ | No concordance effect |
| | Baseline, discordant trials | $31.4 \pm 1.7$ | |
| | Subjective labels (all trials) | $23.3 \pm 0.7$ | |
| | Subjective labels, concordant | $26.3 \pm 1.0$ | Strong concordance effect |
| | Subjective labels, discordant | $17.1 \pm 1.5$ | |
| | Restricted (1 video/emotion) | $50.6 \pm 1.8$ | ↑ despite $1/3$ data |

### Part 1: Intra-subject decoding

The intra-subject baseline (1a) achieved 58.8 ±1.0% accuracy (Fig. 1d), replicating and slightly exceeding the 51.1 ±0.9% reported by Chen et al. (2023).

Restricting the stimulus set from three videos to one per emotion (1b) reduced the data from 90 to 30 seconds of EEG per class, eliminating within-class stimulus variance. Despite this dataset reduction, accuracy increased to 71.3 ±1.3% (chance: 11.1%), compared to 58.8% with three videos (1a).

### Part 2: Cross-subject decoding

The cross-subject baseline (2a) achieved 39.4± 1.1% accuracy (Fig. 2b), replicating and slightly exceeding the 35.2± 1.0% reported by Chen et al. (2023).

Splitting the baseline classifier’s predictions by subjective concordance (2b; Fig. 3a) yielded 33.9 ± 1.4% on concordant trials and 36.0 ± 1.8% on discordant trials. Accuracy was comparable regardless of whether a subject’s dominant self-reported emotion matched the video’s assigned category.

Replacing stimulus-assigned labels with subjective labels (2c; Fig. 3b) reduced accuracy to 26.8 ± 0.9%. Splitting this subjective classifier’s predictions by concordance yielded 31.9 ±1.4% balanced accuracy on concordant trials but only 1.2% on discordant trials (chance: 12.5%).

Restricting the stimulus set to a single video per emotion (2d; Fig. 4c) yielded 44.9± 1.6% accuracy with proper nested cross-validation for hyperparameter selection. This exceeded the baseline accuracy of 39.4% despite using only one-third of the training data.

Replacing LinearSVC with CLISA (2e; Fig. 5, Table 1) reproduced the same confound pattern. The CLISA baseline achieved 34.2± 0.8%. Concordant and discordant trials yielded comparable accuracy (27.7 ±1.1% vs. 31.4 ±1.7%). Subjective labels reduced accuracy to 23.3 ±0.7%, with the concordance split yielding 26.3±1.0% on concordant and 17.1± 1.5% on discordant trials. Restricting to one video per emotion increased accuracy to 50.6 ±1.8%, again exceeding the baseline despite using one-third of the data. A modern deep learning architecture thus does not resolve the stimulus-identity confound.

## Discussion

Our replication yields slightly higher accuracies than those reported by Chen et al. (2023) for LinearSVC (intra-subject: 58.8% vs. 51.1%; cross-subject: 39.4% vs. 35.2%). We used the official pre-computed DE features published with the dataset, whereas the validation code accompanying the dataset re-derives DE features from the pre-processed EEG, which may account for the discrepancy. Our classification code otherwise follows the original methodology. For CLISA, our cross-subject baseline (34.2%) fell below the reported 42.35% despite reusing the original analysis code and matching the pipeline as closely as possible; we were unable to identify the source of this discrepancy.

The intra-subject analyses suggest that above-chance classification reflects autocorrelation confounds rather than shared emotion representations. Two sources of temporal autocorrelation are relevant: intrinsic EEG autocorrelation (temporally adjacent EEG segments have similar spectral content independently of the stimulus) and stimulus-driven autocorrelation (natural movies are temporally autocorrelated, with amplitude spectra following a 1*/f* power law^11,12^). Both allow a classifier to match temporally adjacent train and test segments without learning emotion-related patterns. Restricting to one video per emotion (1b) eliminates within-class stimulus variance and creates a one-to-one mapping between video identity and class label, making both stimulus identity and temporal autocorrelation maximally exploitable. Accuracy increases from 58.8% to 71.3% despite discarding two-thirds of the data, indicating that the additional videos per category diluted the stimulus-identity confound rather than providing learnable emotion signal. Intrinsic EEG autocorrelation and stimulus-driven autocorrelation cannot be disentangled within the present design: increasing the temporal distance between train and test segments within the same video does not isolate intrinsic EEG autocorrelation from stimulus-driven autocorrelation. Both types of autocorrelation are well documented: Xu et al.^13^ recorded EEG from watermelons and showed that labels can be decoded above chance whenever continuous data with the same label is split into training and test sets, even in the complete absence of neural responses (demonstrating intrinsic recording autocorrelation). Kilgallen et al.^9^ identified the repeated-stimulus confound in 16 EEG visual-decoding publications, finding accuracy overestimations of 4.5–7.4%.

The cross-subject analyses paint a consistent picture: the baseline classifier’s performance is driven by stimulus-specific patterns rather than by subjects’ emotional experiences. Concordance analysis (2b) shows that the accuracy is comparable for trials in which subjects reported feeling the labeled emotion (33.9%) and trials in which they did not (36.0%). Replacing stimulus-assigned labels with subjective labels (2c) reduced accuracy from 39.4% to 26.8%, indicating that much of the classifier’s accuracy relies on stimulus-assigned labels rather than subjective reports. The concordance split of 2c provides the most direct test: on discordant trials — where the subject’s felt emotion differs from the stimulus label — balanced accuracy dropped to 15.3%, near the 12.5% chance level. The model trained on subjective labels thus largely fails to predict a subject’s self-reported emotion when it disagrees with the stimulus label, consistent with stimulus decoding rather than emotion decoding. Restricting to one video per emotion (2d) increased accuracy from 39.4% to 44.9% despite using only one-third of the training data, confirming that within-class stimulus variance was noise, not signal, for cross-subject classification as well.

Taken together, these results illustrate violations of three of the six recommendations formulated by Brouwer et al.^7^: the classification is driven by confounding factors rather than by the affective state of interest (R3), the within-video temporal splits violate the independence of training and test data (R4), and the original validation provides no analysis to distinguish stimulus decoding from emotion decoding (R5). R4 is further violated by data leakage in the preprocessing: per-subject z-normalization and LDS smoothing are computed over the entire recording session — including held-out test windows — before the train/test split is applied, creating statistical dependencies between training and test sets^14–16^.

The stimulus-identity confound identified here is not unique to the FACED dataset but likely affects EEG emotion-decoding studies more broadly^17^. Any design with few stimuli per category and within-video train/test splits risks the same confounds. Several recent studies have benchmarked novel deep learning architectures on FACED and reported strong cross-subject accuracies for nine-class “emotion recognition”^18–36^. More critically, FACED has been adopted as the official emotion recognition benchmark in TorchEEG^5^ and Meta’s NeuralBench-EEG v1.0^6^ — our results suggest that what these benchmarks evaluate as emotion recognition may in practice be stimulus recognition.

To illustrate the scope of this problem, consider two of the most widely used EEG emotion datasets. The SEED dataset^2^ uses only 5 film clips per emotion category (∼4 min each) with stimulus-assigned labels — making it susceptible to stimulus confounds. By contrast, the DEAP dataset^3^ is structurally better protected in its original analysis: it uses 40 one-minute music videos, extracts one aggregate feature vector per video, and — crucially — labels each trial with the individual participant’s own subjective rating rather than a stimulus-assigned category. However, many subsequent studies re-analyze DEAP in intra-subject designs by segmenting each video into 1-second windows and performing pooled *k*-fold cross-validation^2,37^, thereby reintroducing the temporal autocorrelation confound. Khan et al.^38^ systematically reviewed 101 DEAP-based studies and found that 87% exhibited at least one such methodological error; when they corrected the most common error, data leakage, classification accuracy dropped from nearly 100% to ∼52%. These examples show that susceptibility arises from specific, identifiable design choices — whether in the experiment itself or in subsequent analyses.

Based on these findings, we propose the following recommendations for future emotion-decoding studies:

1. **Ensure temporal independence between train and test data**. Within-recording train/test splits where train and test segments are temporally adjacent allow classifiers to exploit autocorrelation^9,13^. Blocked designs or leave-one-trial-out schemes where separate stimulus presentations provide independent data are preferable.
2. **Use many distinct stimuli per emotion category**. With sufficiently many stimuli, stimulus identity becomes a nuisance variable that averages out rather than a confound the classifier can exploit.
3. **Use personalized stimuli across subjects**. This mitigates the stimulus-identity confound for cross-subject designs. Personality measures or pre-screening tools can be used to select stimuli likely to elicit the target emotion for each individual, rather than relying on a fixed stimulus set.
4. **Collect per-trial subjective ratings and validate against them**. If a classifier only succeeds with stimulus-assigned labels but fails with subjective labels, it is likely decoding stimuli rather than emotions.
5. **Move beyond declarative ratings**. Multimodal physiological measures — e.g., pupillometry, galvanic skin response, heart rate, respiration — can triangulate emotional states and provide richer ground truth than self-report alone.

Several limitations of the present study should be noted. Declarative self-reports are noisy due to demand characteristics, alexithymia, and task disengagement, and may not reflect true emotional states. Predefined stimulus labels, as used by Chen et al., mitigate this noise but introduce the stimulus-identity confound — the very problem this study identifies. More fundamentally, ground truth for emotions is unavailable; all labels, whether stimulus-assigned or self-reported, are proxies. We focused on the nine-class task; the same design vulnerabilities apply to the binary classification reported by Chen et al.^4^, though with 12 stimuli per class the stimulus identity confound is partially diluted. Finally, we analyzed one dataset and one feature set (DE) with two classifiers (LinearSVC and CLISA). The confound likely generalizes to other methods and datasets with similar designs, but this remains to be demonstrated.

Despite the confounds identified here, the FACED dataset represents a major contribution to the field. It is the largest public EEG emotion dataset (123 subjects), and — crucially — Chen et al.^4^ published both data and code openly. This transparency is what made the present analyses possible: without access to the original features, labels, and classification pipeline, the confounding factors we identify would likely have remained hidden. We commend this practice and urge that it becomes standard. When researchers make their data and code available following FAIR principles^39^, the field gains the ability to self-correct through independent verification and critical re-evaluation — and it is precisely this kind of constructive scrutiny that advances science.

## Author contributions statement

M.G. conceptualized the study, developed the methodology and software, performed the formal analysis and investigation, created the visualizations, and wrote the manuscript. E.S. conducted a systematic review and verification of the analysis code. E.S., A.O., A.H., J.C., D.G., and S.T. reviewed and edited the manuscript.

## Data availability

The FACED dataset was originally published by Chen et al.^4^ and is publicly available on Synapse (https://doi.org/10.7303/syn50614194). We re-used their data for all analyses in this study.

## Code availability

All analysis code is available at

https://github.com/moritz-gerster/faced-stimulus-confound.

## References

1. Li, X. et al. EEG based emotion recognition: A tutorial and review. ACM Comput. Surv. 55, 1–57 (2023).

2. Zheng, W.-L. & Lu, B.-L. Investigating critical frequency bands and channels for EEG-based emotion recognition with deep neural networks. IEEE Trans. Auton. Ment. Dev. 7, 162–175 (2015).

3. Koelstra, S. et al. DEAP: A database for emotion analysis using physiological signals. IEEE Trans. Affect. Comput. 3, 18–31 (2012).

4. Chen, J. et al. A large finer-grained affective computing EEG dataset. Sci. Data 10, 740 (2023).

5. Zhang, Z.Zhong, S.-H. & Liu, Y. TorchEEGEMO: A deep learning toolbox towards EEG-based emotion recognition. Expert. Syst. Appl. 249, 123550 (2024).

6. Banville, H. et al. NeuralBench: A unifying framework to benchmark NeuroAI models. arXiv [cs.LG] (2026).

7. Brouwer, A.-M., Zander, T. O., van Erp, J. B. F., Korteling, J. E. & Bronkhorst, A. W. Using neurophysiological signals that reflect cognitive or affective state: six recommendations to avoid common pitfalls. Front. Neurosci. 9, 136 (2015).

8. Alarcao, S. M. & Fonseca, M. J. Emotions recognition using EEG signals: A survey. IEEE Trans. Affect. Comput. 10, 374–393 (2019).

9. Kilgallen, J. A., Pearlmutter, B. A. & Siskind, J. M. The repeated-stimulus confound in electroencephalography. arXiv [q-bio.NC] (2025).

10. Shen, X., Liu, X., Hu, X., Zhang, D. & Song, S. Contrastive learning of subject-invariant EEG representations for cross-subject emotion recognition. IEEE Trans. Affect. Comput. 14, 2496–2511 (2023).

11. Billock, V. A., de Guzman, G. C. & Scott Kelso, J. A. Fractal time and 1/f spectra in dynamic images and human vision. Phys. D 148, 136–146 (2001).

12. Roberts, M. M., Schira, M. M., Spehar, B. & Isherwood, Z. J. Nature in motion: The tuning of the visual system to the spatiotemporal properties of natural scenes. J. Vis. 22, 7 (2022).

13. Xu, X. et al. Beware of overestimated decoding performance arising from temporal autocorrelations in electroencephalogram signals. arXiv [eess.SP] (2024).

14. Kapoor, S. & Narayanan, A. Leakage and the reproducibility crisis in machine-learning-based science. Patterns (N. Y.) 4, 100804 (2023).

15. Lemm, S., Blankertz, B., Dickhaus, T. & Müller, K.-R. Introduction to machine learning for brain imaging. Neuroimage 56, 387–399 (2011).

16. Mallampati, S. B. & Hari, S. A comparative study on the impacts of data leakage during feature selection using the CIC-IoT 2023 intrusion detection dataset. In 2024 10th International Conference on Electrical Energy Systems (ICEES), 1–6 (IEEE, 2024).

17. Brookshire, G. et al. Data leakage in deep learning studies of translational EEG. Front. Neurosci. 18, 1373515 (2024).

18. Sun, J. et al. Towards recognizing spatial-temporal collaboration of EEG phase brain networks for emotion understanding. In Proceedings of the Thirty-Third International Joint Conference on Artificial Intelligence, 3299–3307 (International Joint Conferences on Artificial Intelligence Organization, California, 2024).

19. Alghamdi, A. M. et al. Cross-subject EEG signals-based emotion recognition using contrastive learning. Sci. Rep. 15, 28295 (2025).

20. Xie, Y., Zheng, Y., Xiao, Z., Lu, W. & Liu, M. Cross-subject EEG emotion recognition based on temporal asynchronous alignment contrastive learning. arXiv [cs.HC] (2026).

21. Wang, F.Tian, Y.-C. & Zhou, X. Cross-dataset EEG emotion recognition based on pre-trained vision transformer considering emotional sensitivity diversity. Expert. Syst. Appl. 279, 127348 (2025).

22. El Ouahidi, Y. et al. REVE: A foundation model for EEG – adapting to any setup with large-scale pretraining on 25,000 subjects. arXiv [cs.LG] (2025).

23. Ding, Y. et al. EmT: A novel transformer for generalized cross-subject EEG emotion recognition. IEEE Trans. Neural Netw. Learn. Syst. 36, 10381–10393 (2025).

24. Gao, D. et al. A multi-domain constraint learning system inspired by adaptive cognitive graphs for emotion recognition. Neural Netw. 188, 107457 (2025).

25. Xiao, Q. et al. BrainOmni: A brain foundation model for unified EEG and MEG signals. In The Thirty-ninth Annual Conference on Neural Information Processing Systems (2025).

26. Ye, W. et al. Semi-supervised dual-stream self-attentive adversarial graph contrastive learning for cross-subject EEG-based emotion recognition. IEEE Trans. Affect. Comput. 16, 1–16 (2024).

27. Li, D., Huang, S., Xie, L., Wang, Z. & Xu, J. Neuron perception inspired EEG emotion recognition with parallel contrastive learning. IEEE Trans. Neural Netw. Learn. Syst. 36, 14049–14062 (2025).

28. Wang, J. et al. EEGMamba: An EEG foundation model with mamba. Neural Netw. 192, 107816 (2025).

29. Li, W., Fan, L., Shao, S. & Song, A. Generalized contrastive partial label learning for cross-subject EEG-based emotion recognition. IEEE Trans. Instrum. Meas. 73, 1–11 (2024).

30. Zhou, Y. et al. SPICED: A synaptic homeostasis-inspired framework for unsupervised continual EEG decoding. In The Thirty-ninth Annual Conference on Neural Information Processing Systems (2025).

31. Zhang, H., Zuo, T., Chen, Z., Wang, X. & Sun, P. Z. H. Evolutionary ensemble learning for EEG-based cross-subject emotion recognition. IEEE J. Biomed. Health Inform. 28, 3872–3881 (2024).

32. Chang, J., Zhang, Z., Qian, Y. & Lin, P. Multi-scale hyperbolic contrastive learning for cross-subject EEG emotion recognition. IEEE Trans. Affect. Comput. 16, 1716–1731 (2025).

33. Li, C., Wang, F., Zhao, Z., Wang, H. & Schuller, B. W. Attention-based temporal graph representation learning for EEG-based emotion recognition. IEEE J. Biomed. Health Inform. 28, 5755–5767 (2024).

34. Ma, J. et al. CodeBrain: Bridging decoupled tokenizer and multi-scale architecture for EEG foundation model. In The Fourteenth International Conference on Learning Representations (2026).

35. Hu, M., Xu, D., He, K., Zhao, K. & Zhang, H. Cross-subject emotion recognition with contrastive learning based on EEG signal correlations. Biomed. Signal Process. Control. 104, 107511 (2025).

36. Chen, H. et al. VAE-CapsNet: A common emotion information extractor for cross-subject emotion recognition. Knowl. Based Syst. 311, 113018 (2025).

37. Zhu, Y. & Zhong, Q. Differential entropy feature signal extraction based on activation mode and its recognition in convolutional gated recurrent unit network. Front. Phys. 8 (2021).

38. Khan, N. N. et al. The role of review process failures in affective state estimation: An empirical investigation of DEAP dataset. arXiv [eess.SP] (2025).

39. Wilkinson, M. D. et al. The FAIR guiding principles for scientific data management and stewardship. Sci. Data 3, 160018 (2016).

